# Increased influence of periphery on central visual processing in humans during walking

**DOI:** 10.1101/382093

**Authors:** Liyu Cao, Barbara Händel

## Abstract

Cognitive processes are almost exclusively investigated under highly controlled settings while voluntary body movements are suppressed. However, recent animal work suggests differences in sensory processing between movement states by showing drastically changed neural responses in early visual areas between locomotion and stillness. Does locomotion also modulate visual cortical activity in humans and what are its perceptual consequences? Here, we present converging neurophysiological and behavioural evidence that walking leads to an increased influence of peripheral stimuli on central visual input. This modulation of visual processing due to walking is encompassed by a change in alpha oscillations, which is suggestive of an attentional shift to the periphery during walking. Overall, our study shows that strategies of sensory information processing can differ between movement states. This finding further demonstrates that a comprehensive understanding of human perception and cognition critically depends on the consideration of natural behaviour.

## Introduction

Perception is not only a function of the stimulus, but also very much influenced by internal factors such as arousal and attention. Recently, animal work has added an interesting dimension to this: the behavioural state such as locomotion. In mice (1-9) as well as invertebrates (10-12), these studies have shown a drastic difference in neural responses in sensory areas between locomotion and still state that goes beyond the influence of arousal (for a recent review see (13, 14)). Unfortunately, our understanding of human visual cognition in natural settings such as during walking is surprisingly limited, and whether the influence of locomotion on early sensory activity translates to humans is unclear. Behavioral studies (15) and some emerging electrophysiological work indicate a link between movement and cognition such as memory, attention (16-18) and perceptual processes (19-21) in humans. However, due to technical constraints, research focused on the study of humans in stationary settings, such as walking on a treadmill or stationary cycling (17, 22, 23), with few exceptions (24-26). In the current study, we combine the latest mobile EEG/EOG technology, mobile visual stimulation and behavioural measurements in freely walking humans. By using steady state visual evoked potentials (SSVEP), which are known to originate from early visual cortex (27), and contrast modulated surround suppression effects, we demonstrated that walking modulates sensory processing in early visual cortex, which is in line with work on freely moving animals (28). Moreover, we show that this modulation is linked to alpha oscillations, indicating an attention-like process.

## Results and Discussion

Participants were asked to stand still, walk slowly or walk with normal speed while completing a perceptual task, which was presented via a head-mounted display (Fig. 1A). Participants fixated a centrally presented circular grating flickering at 15 Hz (Fig. 1B). Flicker introduces a strong entrained steady state visually evoked potential, with a focus in early visual areas (27). The task was to detect a threshold titrated contrast change (target) presented randomly in time and location within the flickering central grating. A stable background grating surrounded the central grating showing one of four different contrast levels between 0 and 100% (Fig. 1B). Increased surround contrast has been previously reported to inhibit responses to the central input, which is termed surround suppression (29). For further analysis, the 15Hz SSVEP power values were obtained in a 2-second time window prior to target onset. SSVEP was readily detected during walking (as reported by previous studies testing the signal quality of mobile EEG setups (22, 30)) and showed an occipital focus (Fig. 1C). SSVEP power values were analyzed further as a function of the two experimental manipulations, i.e. walking condition and surround contrast level.

**Fig. 1.**
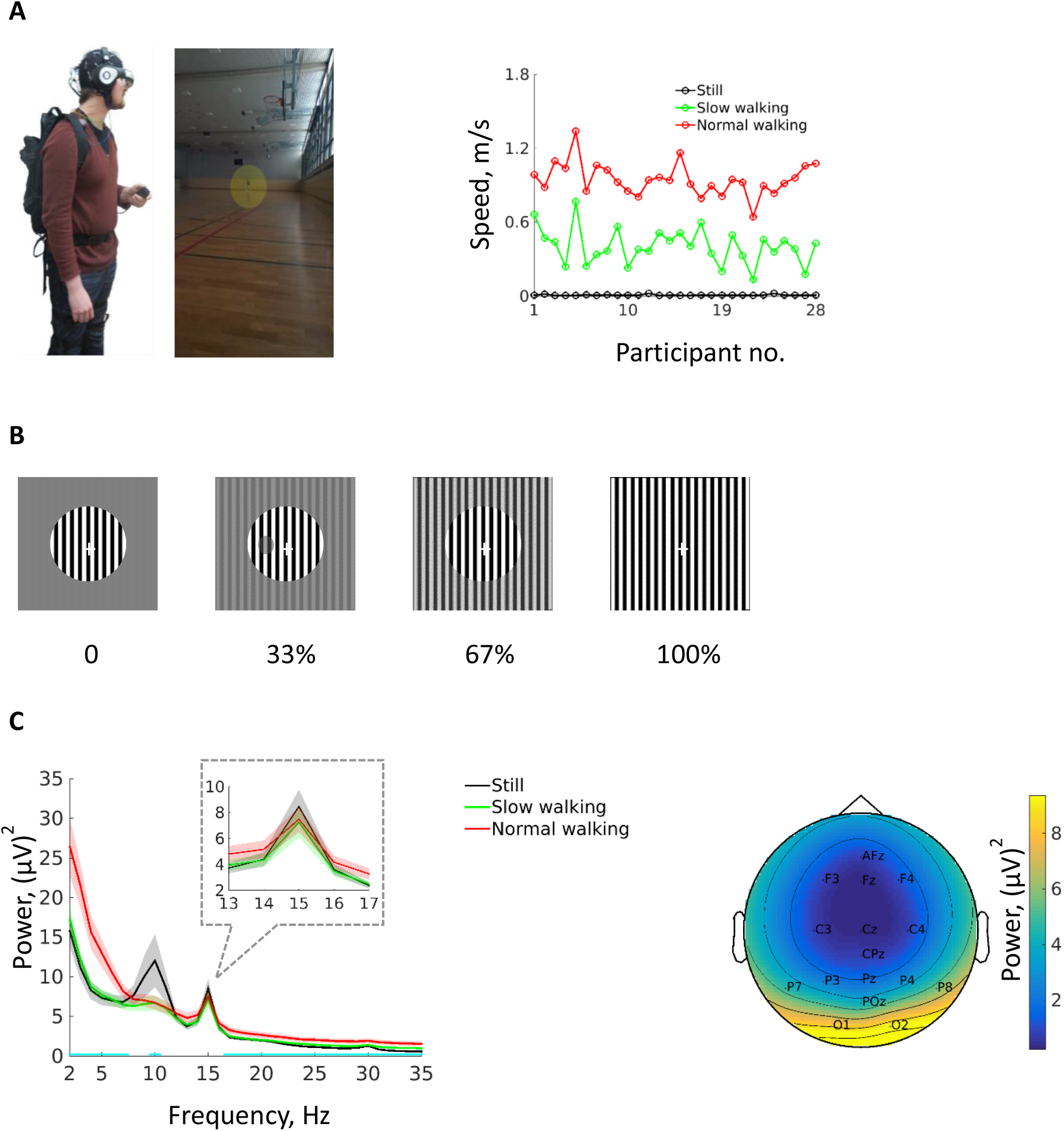
Experimental set-up and SSVEP responses. (A) Participants carried all experimental equipment and walked freely in a 45×27 metre sports hall. Walking speed is shown on the right (data from two participants are missing due to technical errors during the recording). (B) Illustration of the central flickering grating (visual angle: 6.1°) and the four different levels of surround contrast (visual angle: 21.5°). An example contrast change (target) is shown in the second row (see the methods section for correct stimuli scales). (C) Power spectrum of EEG responses in different walking conditions. Inset shows power responses close to the SSVEP frequency of 15 Hz. Cyan lines mark the frequencies that showed power differences between walking conditions (*p* < 0.05, no corrections for multiple comparisons). Shading indicates ±1 standard error, n = 25 participants. Scalp topography of the SSVEP response is shown on the right.

### Influence of Surround Contrast: EEG Data

A within-subjects 2-factorial (3×4) ANOVA (F1: walking condition; F2: contrast level) revealed significant main effects (correction with Greenhouse-Geisser where necessary) (Fig. 2 A and B). As expected (31), SSVEP power decreased with increasing contrast level (F(3,72) = 5.32, *p* = 0.02). Increasing walking speed also was associated with significantly decreased SSVEP amplitudes (F(2,48) = 9.31, *p* = 0.002). Most interestingly, there was a significant interaction between walking condition and contrast level (F(6,144) = 3.51, *p* = 0.01). Post-hoc analysis showed that SSVEP power was significantly decreased due to increased contrast level in the two walking conditions but not during standing still, i.e. significant surround suppression was only observed in walking conditions (for detailed statistics see SI, Table S1). We excluded the possibility that the increased influence of the surround contrast during walking was caused by a shift of gaze or attention towards the lower visual field, i.e. near the feet, by showing that the bottom target did not have increased behavioural or electrophysiological relevance as compared to other targets (SI, Fig S1, S5). Importantly, the interaction effect between walking condition and contrast level on SSVEP power was preserved during target presentation (200 – 600 ms after the target onset) (SI, Fig. S1). In sum, we demonstrated that walking can modulate visual-related signal processing in the brain.

**Fig. 2.**
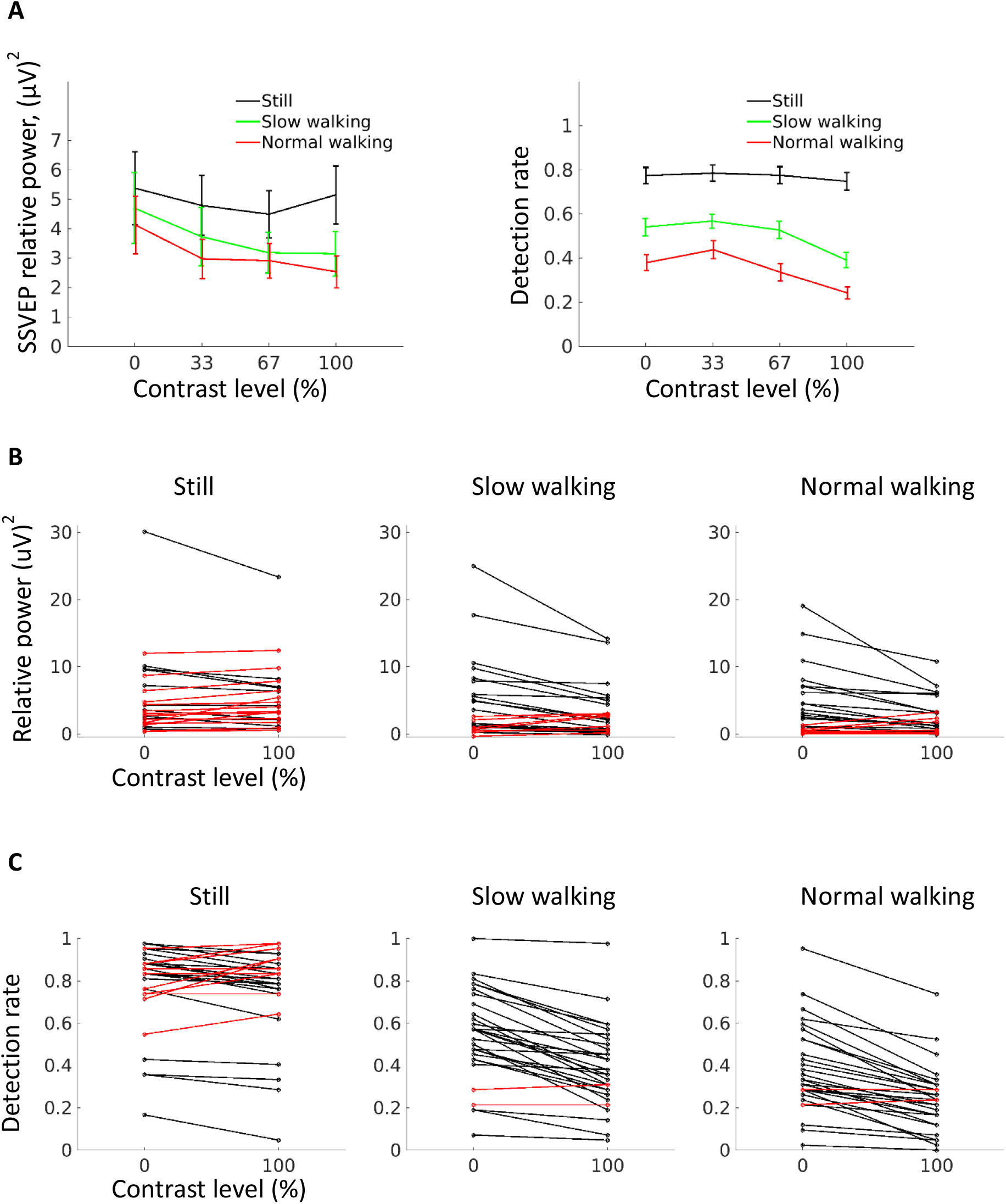
Modulation of SSVEP responses and behavioural responses by surround contrast in different walking conditions. (A) (left) SSVEP responses significantly decreased from 0 surround contrast level to the other three levels of contrast in both walking conditions, but not in the standing still condition. Vertical lines indicate ±1 standard error, n = 25 participants. The detection rate of the target showed a similar pattern as SSVEP (right). Vertical lines indicate ±1 standard error, n = 30 participants. Refer to SI Appendix, Table S1 for break-down statistics. (B) Individual data from contrast level 0 and 100% for SSVEP. Red lines indicate participants whose data patterns do not conform to surround suppression. (C). Individual data from contrast level 0 and 100% for detection rate. Red lines indicate participants whose data patterns do not conform to surround suppression.

Given the great importance of any possible role of movement artefacts in data recorded in freely moving participants, artefacts will be discussed at length in the supplementary information. However, we want to shortly resolve three main points of concern: 1. Could our signal picked up with EEG during walking be muscle activity? Since we compared the power of SSVEP, which is known to originate from early visual cortex, we can be confident that the signal we analysed is not dependent on artefacts. We additionally checked for the interaction effect in other frequencies by repeating the ANOVA from 3 to 30 Hz (step size 1). No other frequencies had *p* values lower than 0.05, except the signal of 17 Hz which took contribution from 15 Hz signal (signals from each frequency was referenced to the mean of 4 nearby frequencies as was for SSVEP signal; Fig. S1). This further shows that our effect is frequency specific and not due to a shift in overall power. 2. Different walking conditions lead to different signal to noise ratios, which can lead to a difference in detected activity between walking conditions. Could this explain our observed effects? While the overall power of SSVEP between walking conditions might be influenced by a changed signal to noise ratio, our approach to compare power only between surround contrast conditions and not between walking conditions circumvents this problem. 3. Eye movements differ between walking conditions and therefore lead to a difference in visual input. Could this cause a contrast dependent change in the SSVEP? In general, the above given argument holds here as well. We did not compare walking conditions, therefore only if eye movements would be modulated by surround contrast differently for different walking speed, eye movement effects could explain our data. We measured EOG with verified spatial sensitivity to saccades as small as 0.1° (SI, Fig. S3). Careful analysis revealed that there was a modulation of blink and saccade rate for different walking conditions (SI, Fig. S3, S4). However, no interaction with surround contrast could be found. Additionally, when only including trials without saccades or blinks in our main analysis, a similar pattern of SSVEP interaction between walking condition and surround contrast was observed (SI, Fig. S3, S4).

### Influence of Surround Contrast: Behavioural Data

Our physiological data are paralleled by behavioural results (Fig. 2 A and C), which further reduces the likelihood of an artefact-based effect. The same within-subjects 2-factorial (3×4) ANOVA (F1: walking condition; F2: contrast level) revealed significant main effects of walking condition (F(2,58) =107.12, *p* < 0.001) and contrast level (F(3,87) = 54.91, *p* < 0.001) on detection rate. Detection rate decreased with increasing walking speed and contrast levels (see also SI for discussion. Research suggests that walking cannot be fully automatic (32) and can impair spatial memory capacity and target detection time (33). The overall negative effect of walking on detection rate might be explained by the dual task demands during walking.

A significant interaction (F(6,174) = 9.35, *p* < 0.001) indicated that, in line with the neurophysiological finding, a perceptual surround suppression effect was present during walking only (for detailed statistics see SI, Table S1). There is a slight difference in the modulation curve between behavioural and electrophysiological measures. This could be explained by the fact that the detection rate is critically dependent on the surround contrast as well as the target threshold. The SSVEP strength, however, is only modulated by the surround contrast. See SI for details.

No interaction effects between walking and surround contrast were found for reaction time or false alarms (SI, Fig. S2), suggesting that it is not a motor based effect such as lowering the motor response threshold. Please note that it was not the overall lower detection rate during walking that led to a non-linear influence of surround contrast. When forcing the detection rate for the 0% surround contrast to be similar for normal walking and standing still, we still found a greater influence of surround contrast during walking (SI, Fig. S2).

### Interaction Between SSVEP Power and Alpha Power

Under static condition, surround suppression increases with attention focused on the surround (34). We therefore assume that walking does not lead to a general drop in attentional resources but a walking-induced specific shift of attention towards the periphery, thereby leading to the observed increased influence of surround contrast. This interpretation is further supported by our finding that alpha power, a ~10 Hz oscillation closely associated with attention, co-varies with the observed SSVEP power.

The difference in alpha power between walking conditions is depicted in Fig. 1B. A general decrease in alpha due to walking has been noted previously (22, 35), but a functional role was not experimentally tested. One influential account of alpha oscillations from neurocognitive studies proposes that alpha acts to functionally block out irrelevant information (36). High alpha power is thereby associated with inhibiting sensory processing in a locally specific fashion (37). This means that if attention is directed to the fovea, alpha can inhibit peripheral input processing and vice versa. To test the link between alpha power and suppression effects due to peripheral input, trials were sorted according to the strength of SSVEP power and divided into weak SSVEP and strong SSVEP trial groups based on the median. This analysis was performed separately for each walking condition. A significant difference in the power of alpha band was observed for all three walking conditions (standing still: t(24) = −2.78, *p* = 0.01; slow walking: t(24) = −3.87, *p* < 0.001; normal walking: t(24) = −3.91, *p* < 0.001). Strong alpha power was associated with strong SSVEP power (Fig. 3). No significant differences in other frequency bands were found. To corroborate the link between alpha power and SSVEP power, we performed a multiple linear regression analysis on a single trial basis taking the SSVEP as the dependent variable, alpha power and number of saccades and blinks as predictors. Results showed that alpha power positively predicted SSVEP power (t(24) = 4.82, *p* < 0.001), in addition to the negative prediction effects from both saccades (t(24) = −2.86, *p* = 0.009) and blinks (t(24) = −5.48, *p* < 0.001).

**Fig. 3.**
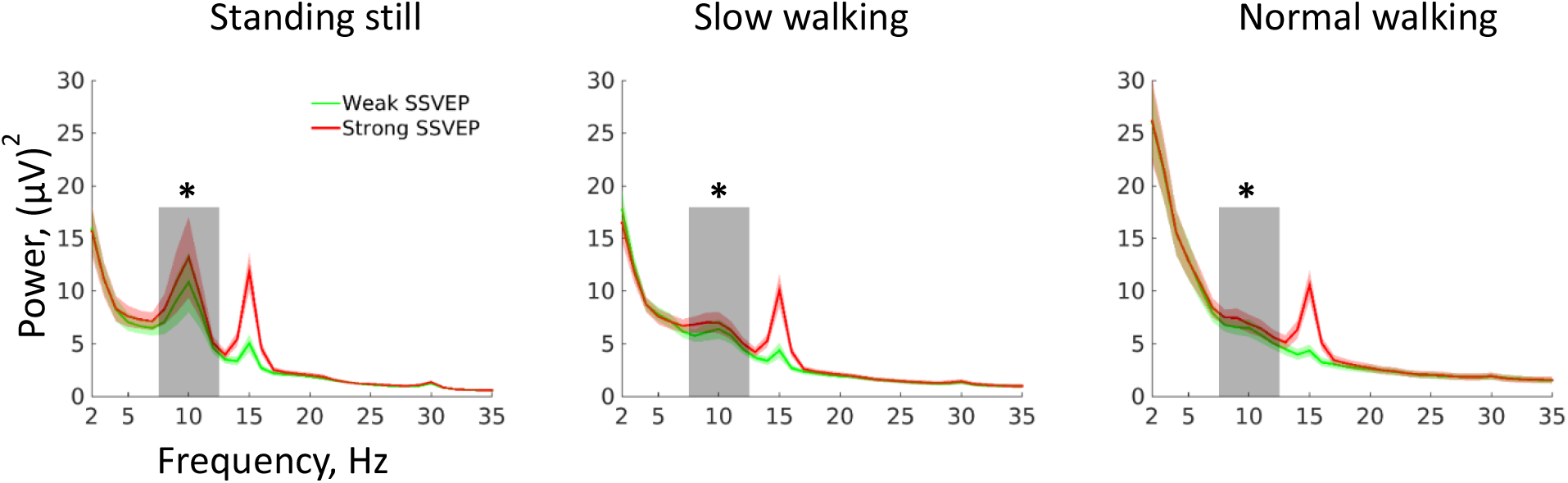
Covariation between SSVEP power and alpha power. In all the three walking conditions, stronger SSVEP power was associated with stronger alpha power. No significant differences in other frequency bands were found. Shading indicates ±1 standard error. Grey bars indicate the alpha band (8 – 12 Hz) and asterisks indicate significant differences, n = 25 participants.

Unfortunately, our low number of electrodes does not allow for a meaningful source localization of alpha power. We therefore cannot conclusively argue that the observed decrease in alpha due to walking indeed originates from areas within V1 dealing with peripheral vision. Nevertheless, our data are consistent with the idea that while standing still, high alpha power may help blocking out influence from the periphery thus attenuating surround suppression. During walking, however, peripheral alpha power is decreased, thus peripheral blocking is compromised, and the surround contrast can exert its suppressive impact on the central grating. The alpha decrease during walking is therefore associated with an increased SSVEP power.

It is biologically plausible that walking enhances peripheral processing since peripheral input holds important cues for locomotion and navigation (38, 39). The observed pattern of decreased alpha power during walking could constitute an attentional mechanism to promote peripheral input during walking. Despite slightly different findings in mice, the above suggestion follows an argument based on animal work (28), stating that walking affects surround suppression to an extent comparable to attention. The increase in surround suppression during walking therefore falls in line with recently reported effects of attention on surround suppression in monkey V4 (34). We demonstrate, for the first time, an impact of free walking on surround suppression in humans which escaped clear detection during restricted walking on a treadmill (40). One reason might be that walking on a treadmill is biomechanically different from overground walking (41) and navigation and accompanying sensory processes are unnecessary if locomotion is restricted to one place. Our approach of free walking may be extended by applying recent exciting technological developments (42).

Overall, we propose that the processing of sensory information crucially depends on the movement of the subject and further emphasize the call for natural settings when investigating perceptual mechanisms and cognitive processes in general.

## Methods

### Participants

30 healthy participants (20 females; mean age: 29.2; STD: 8.0) were recruited from a local participant pool. All participants gave written informed consent prior to the study and received monetary compensation after the study. The study was approved by the local ethics committee (Department of Psychology, University Würzburg) and was conducted in accordance with the Declaration of Helsinki and the European data protection law (GDPR).

### Stimuli and Task

A circular grating (visual angle: 6.1°; spatial frequency: 0.05 cycle/pixel) flickering at 15 Hz between 100% contrast and 0 contrast was presented at the centre of the screen (refreshing rate: 60 Hz; resolution: 1280 x 720). The contrast level was defined as deviations from plain grey, which had a luminance level of 35 cd/m^2^ (0 contrast means plain grey and 100% contrast means alternations between plain white and plain black). The central grating was surrounded by a fullscreen surround contrast (visual angle: 21.5°; spatial frequency: 0.05 cycle/pixel; in phase with the central grating), which had one of the four following contrast levels: 0, 33%, 67%, and 100%. The behavioural task was to detect a briefly presented disk contrast target (duration: 500 ms; visual angle: 1.2°; deviation from the screen centre: 2.0°) within the central grating by pressing a button as quickly as possible independent of the location of the target. The threshold of the target was determined using a 1-up-4-down procedure with 0 surround contrast(43). Participants completed the threshold test (about 5 minutes) while sitting down prior to the formal testing. The target appeared randomly in one of four possible locations: above, below, to the left or to the right of a central cross (visual angle: 0.3°), on which participants were required to keep fixation throughout the testing period. Stimuli were created and controlled with Psychtoolbox-3 on Matlab (The MathWorks Inc., USA).

Participants then completed the main task in three different walking conditions: standing still, walking slowly or walking with a normal speed. There were 7 testing blocks for each speed (21 blocks in total; random order). Before the start of each block, participants received instruction about the walking condition during the block. In each block, each of the four different surround contrast levels were presented for 35 seconds (random order). During each 35-second period, 6 targets were presented in the interval of 5 – 31.5 second with a stimulus onset asynchrony randomly sampled between 3.5 and 6.5 seconds. After each block, participants were encouraged to take a break. A technical stop was made after block 7 and block 14, during which the EEG electrode impedance was checked. After finishing the test, we also collected EEG data of free walking in a task-free setting during light or darkness and some questionnaires were completed. Additionally, EEG data were recorded while subjects executed saccades between 0.1-5degrees by following a saccade target presented on an external screen. Data concerning these controlled saccades are presented in the supplementary information. The study was conducted in an activity hall (about 30 m x 50 m; wooden floor) of the university gym. Since the lower rim of the visual field was unobstructed, participants could freely move around the large testing field using the information required for navigation (44) from the lower visual field.

### Equipment

A Dell laptop (model: Latitude E7440) was used for running the experiment program and for data collection. During the experiment, the laptop was put into a rucksack which was carried by participants to ensure that participants were fully mobile during the testing. The screen of the laptop, which was used for stimulus presentation, was projected to through a video headset (Glyph Founder’s edition, Avegant Corporation, USA). Behavioural responses were collected using a hand-held response button (model: K-RB1-4; The Black Box ToolKit Ltd, UK), which is connected to the laptop with a USB connection. EEG data were collected using a Smarting mobile EEG system (mBrainTrain LLC, Serbia), which has 24 recording channels with a sampling rate of 250 Hz. We used 6 channels for EOG recording (for each eye: one below and one above the eye, one to the outer canthus), 2 channels for possible re-referencing (attached to both earlobes; eventually not used for the study), and the remaining 16 channels for EEG recording (see Fig. 1C for EEG channel distribution). A common mode sense active electrode placed between Fz and Cz was used for online reference. The EEG signal amplifier and data transmitter are integrated into a little box (82 x 51 x 12 mm; 60 grams) which is attached to the back of the EEG cap. Data transmission is achieved via Bluetooth. The EEG system also has a build-in gyroscope, which was used to measure head movements. Motion data (speed and acceleration; sampling rate: 120 Hz) were collected using a Perception Neuron system (Noitom Ltd, China). Three functional motion sensors were attached to participants’ back, left and right foot ankles. Triggers for recording stimulus timing and behavioural responses were generated with the software Lab Streaming Layer (https://github.com/sccn/labstreaminglayer), which was also used for collecting and synchronizing other streams of data (EEG and motion data). See Fig. 1A for an illustration of the set-up.

### Data Analysis

Data analysis was performed with the Fieldtrip toolbox(45) and in-house scripts. The within-subjects ANOVA was performed using SPSS-22 (IBM Corp., USA) (Greenhouse-Geisser correction was performed when the sphericity assumption was violated). Throughout the manuscript, statistical cut-off was taken at the *p* value of 0.05 and all t-tests are two-tailed.

EEG data from 5 participants were incomplete due to data transmission error, therefore only the remaining 25 full EEG datasets were included for the following analysis. For each participant, continuous EEG data were first cut into 35-second epochs, which covers the full duration of each surround contrast level (84 epochs per participant: 3 speeds x 7 blocks x 4 surround contrast levels). The data were then high-pass filtered at 1 Hz (a windowed sinc finite-impulse-response filter with Kaiser windowing was used for all the filtering processes unless otherwise stated) and low-pass filtered at 100 Hz. Noisy epochs (identified through visual inspection) were repaired through interpolation. 124 epochs in total (from 6 participants) were repaired. Among them, 8 epochs (from 3 participants) were included in the SSVEP analysis (see below). Next, data were further segmented into a 2-second snippet ending at the target onset time point. All the analyses below were based on these 2-second trials that are free from stimulus changes (i.e. target onset) and button press responses (trials with a button press in the 2.5-second window before the target onset were excluded). On average, 496.0 trials (STD: 11.0) remained for each participant (504 trials before rejection).

EEG power spectrum was obtained using the Welch’s method. For each participant, 3 channels from the occipital area with highest mean power of all trials at 15 Hz were selected. A further trial exclusion was performed within each speed condition based on the high-frequency power using the MAD-median rule: let p be the sum of the high-frequency power (20 – 99 Hz) of a single trial and P be the high-frequency power of all trials within a speed condition. If |p – median(P)| x 0.6745 > 2.24 x MAD (the median absolute deviation from the median), this trial is an outlier(46). On average, 450.5 trials (STD: 24.7) remained after this step.

A within-subjects ANOVA was performed to compare the power of each frequency (between 2 and 35 Hz in steps of 1 Hz) among the three walking conditions (Fig. 1B). No corrections for multiple comparison were made as this analysis was conducted to give a first overview of the data. Conclusions are based on the statistical approach as described below. The SSVEP power at 15 Hz was then referenced to the mean power of nearby frequencies (13, 14, 16, and 17 Hz) through subtraction. The referenced SSVEP power was averaged within each speed/surround contrast combination before being subjected to a 3 (walking condition) by 4 (contrast level) within-subjects ANOVA.

To find the association between SSVEP power and alpha power, trials were grouped based on the median raw SSVEP power into weak SSVEP and strong SSVEP trials separately for each speed condition. For both walking conditions, 5 trials with lowest high-frequency power in the weak SSVEP group and 5 trials with highest high-frequency power from the strong SSVEP group were excluded. This eliminated the group difference in the high frequency band but did not change the statistical results comparing group differences in other frequency bands. Possible power difference in the delta band (2 – 3 Hz), theta band (4 – 7 Hz), alpha band (8 – 12 Hz), and high-frequency band (20 – 99 Hz) was compared using within-subjects t-tests across participants. Furthermore, a multiple linear regression analysis was performed for each participant taking the SSVEP power of a single trial as the response variable, alpha power (from individual peak alpha frequency) and numbers of blinks and saccades as predictors. Both SSVEP power and alpha power were normalized within each testing block (i.e. fixed speed) to account for problems of power shifts between different speed conditions. Statistics was performed by making a group-level t-test between the slope parameter and 0.

### Behavioural Responses

A hit response was recorded if a button press was made within 1 second from the onset of the target. All other responses during the testing were regarded as false alarms. A detection rate was calculated for each speed/surround contrast combination before being subjected to a 3 (walking condition) by 4 (contrast level) within-subjects ANOVA. Data from all 30 participants were included for analysis. False alarms and Reaction time data (5 participants were excluded based on the criterion that less than 5 responses were made in at least one condition) were analysed similarly. Note that arcsine transformed detection rate data and square root transformed data for false alarms resulted in similar statistics results. Therefore, no data transformation was performed for the reported statistics.

To test if increased surround suppression during walking is simply due to the overall decrease of detection rate, we focused our analysis on the target (selected individually for each participant) with lowest detection rate in standing still condition and the target with the highest detection rate in normal walking condition. We further excluded the first eight participants in the rank of detection rate difference under 0 surround contrast between standing still condition and normal walking condition, after which a comparable detection rate under 0 surround contrast between standing still and normal walking was achieved for the remaining 22 participants. Comparisons of detection rate were then continued for the other 3 contrast levels (SI, Fig. S2).

### Blink Detection

Blinks were detected from the vertical EOG component, i.e. the amplitude difference between the EOG channels above and below eyes. The vertical EOG component was high-pass filtered at 0.2 Hz and low-pass filtered at 20 Hz. A blink was detected if the vertical component crossed a threshold of 20 μV. Blinks with peak amplitude lower than 40 μV or amplitude standard deviation smaller than 15 μV were excluded. Adjacent blink points within 100 ms were combined into one blink. Results of blink detection from both eyes were quite similar, therefore only results from the left eye were used.

### Saccade Detection

Saccade detection was based on the so-called REOG component, which is the difference between the mean of all 6 EOG channels and the Pz channel. REOG component was band-pass filtered between 20 and 90 Hz using a 6th order Butterworth filter and then a Hilbert transform was performed to obtain the amplitude envelop. All data points where the amplitude value deviated from the mean by 2.5 standard deviations were considered saccade-related and were grouped into one saccade if they were within 20 ms apart. This was done separately for each of the 21 recording blocks. This method was shown to be able to detect saccades very reliably (47).

### Strong Head Movements Detection

Strong head movements were detected from the gyroscope data. Time points with the rotational velocity deviating from the mean deviations (calculated within each testing block) by 2.5 standard are regarded as strong head movement points. Strong head movement points within a 2-second window were assigned to one head movement event.

## Data availability

The data and Matlab codes are available from the corresponding author upon reasonable request.

## Acknowledgements

We would like to thank Reinhard Roth for providing access to the facilities at the sports centre of the University of Würzburg, Wilfried Kunde for his support, and Anne Boeckler-Raettig and Marieke Schoelvinck for helpful comments. We further thank Normann Mangold for technical support, the mBrainTrain team for support on the mobile EEG setup and Stefan Debener for technical advice in the planning phase. This study was supported by a starting grant from the European Research Council awarded to BH (grant number 677819).

## Author Contributions

L.C. implemented the testing protocol, collected, and analyzed the data; B.H. developed the experiment and led the project; L.C. and B.H. wrote the manuscript.

